# Antitumor activity of a lectibody targeting cancer-associated high-mannose glycans

**DOI:** 10.1101/2021.04.28.441869

**Authors:** Young Jun Oh, Matthew W. Dent, Angela R. Freels, Qingwen Zhou, Carlito B. Lebrilla, Michael L. Merchant, Nobuyuki Matoba

## Abstract

Aberrant protein glycosylation is a hallmark of cancer, but few drugs targeting cancer glycobiomarkers are currently available. Here, we show that a “lectibody” consisting of the high-mannose glycan-binding lectin Avaren and human IgG1 Fc (AvFc) selectively recognizes a range of cell lines derived from lung, breast, colon and blood cancers at nanomolar concentrations. AvFc’s binding to the non-small cell lung cancer (NSCLC) cell lines A549 and H460 was characterized in detail. Co-immunoprecipitation proteomics analysis revealed that epidermal growth factor receptor (EGFR) and insulin-like growth factor 1 receptor (IGF1R) are among the lectibody’s common targets in these cells. AvFc blocked the activation of EGFR and IGF1R by their respective ligands in A549 cells and inhibited the migration of A549 and H460 cells upon stimulation with EGF and IGF1. Furthermore, AvFc induced potent Fc-mediated cytotoxic effects and significantly retarded A549 and H460 tumor growth in SCID mice. Immunohistochemistry analysis of primary lung tissues from NSCLC patients demonstrated that AvFc preferentially binds to tumors over adjacent non-tumor tissues. Our findings provide evidence that increased abundance of high-mannose glycans in the glycocalyx of cancer cells can be a druggable target, and AvFc may provide a new tool to probe and target this tumor-associated glycobiomarker.

## INTRODUCTION

It has become evident that changes in protein glycosylation patterns are associated with various disease conditions, including viral infections and cancer.^1, 2^ One such change observed in several cancer types is a significant increase in the proportion of high-mannose-type glycans, which constitute a type of asparagine-linked glycan (*N*-glycan) containing 5-9 terminal mannose residues.^3, 4^ In normal cells, these glycoforms appear in the endoplasmic reticulum (ER) but are subsequently processed into complex-type glycans by a series of mannosidases and glycosyltransferases in the Golgi apparatus as nascent glycoproteins passage through the secretory pathway. Thus, high-mannose glycans are considered to be “immature” *N*-glycans that are generally confined in the ER under normal conditions.^1^ However, recent studies based on quantitative mass spectrometry analyses of cancer tissue have demonstrated that this may not always be the case. For example, high-mannose glycans were elevated in serum samples from breast cancer patients, which correlated with cancer progression.^5^ Analysis of large cohorts of paired breast cancerous and adjacent non-tumor tissues found a high-mannose glycan (Man8) along with a triantennary glycan to be dramatically increased in the membrane fraction of tumors.^6^ Increased abundance of high-mannose glycans has also been observed in colorectal tumor tissues^7-9^, hepatocellular carcinoma^10, 11^, metastatic cholangiocarcinoma^12^, lung adenocarcinoma^13^, pancreatic cancer^14^, ovarian cancer^15, 16^, prostate cancer^17^ and skin basal cell carcinoma and squamous cell carcinoma^18^. Collectively, the aberrant increase of high-mannose glycans on malignant cells may provide a unique biomarker for drug development. Nevertheless, there are few agents that can distinguish tumor-associated high-mannose glycans from other glycoforms present on a normal cell’s surface, and thus their druggability remains unclear.

Previously, we reported the creation of an antibody-like “lectibody” molecule comprised of the oligomannose-specific Avaren lectin and the fragment crystallizable region (Fc) of human IgG1, called Avaren-Fc (AvFc).^19^ Avaren is an engineered variant of the actinomycete-derived antiviral lectin actinohivin,^20, 21^ with amino acid substitutions to improve solubility and producibility. AvFc neutralized the infectivity of multiple HIV strains and hepatitis C viruses at nanomolar concentrations through high-affinity binding to high-mannose glycans clustered on their envelope glycoproteins.^19, 22^ Additionally, the lectibody exhibited antibody-dependent cell-mediated virus inhibition against HIV-infected peripheral blood mononuclear cells (PBMCs) via its capacity to interact with activating Fcγ receptors such as FcγRI and FcγRIIIa. Preliminary safety studies in mice and rhesus macaques showed that systemic administration of AvFc did not induce any discernable toxicity.^19^ Furthermore, systemic administration of 25 mg/kg of AvFc every other day (Q2D) for 14 or 20 days (8 or 11 doses total, respectively) completely protected against HCV challenge without causing hepatotoxicity or any other significant adverse effects in a chimeric human liver mouse model.^22^ These results lend support for the use of AvFc in novel therapeutic strategies targeting high-mannose glycans that may loom on the cell surface in high densities under pathological conditions.

In light of growing evidence for the abberant overexpression of high-mannose glycans in neoplastic cells, we hypothesized that AvFc could efficiently recognize these mannose-rich glycans on the surface of cancer cells and thereby exhibit antitumor activity. To address this hypothesis, the present study investigated the lectibody’s capacity to target cancer using human non-small cell lung cancer (NSCLC) cell lines, murine xenograft models of human NSCLC and primary human NSCLC tissue sections. Our results provide implications for a novel anti-cancer strategy targeting tumor-associated high-mannose glycans.

## RESULTS

### AvFc selectively recognizes various cancer cell lines

Given that high-mannose glycans are elevated in various neoplastic cells and tissues,^5-8, 10, 13-17^ we first tested whether AvFc can effectively recognize cancer cells. Flow cytometry analysis showed that the lectibody bound to a range of human cancer cell lines derived from breast, lung, colon, blood, cervical, and prostate tumors at nanomolar concentrations. Nanomolar concentrations (0.1 – 10 µg/mL) of AvFc exhibited distinct binding to most of the 27 cancer cell lines tested, albeit with varying degrees of efficiency. MDA-MB-231 breast carcinoma, A549 lung adenocarcinoma, H460 large-cell lung carcinoma, HT-29 colon adenocarcinoma, SK-MEL-2 melanoma, and HeLa cervical carcinoma cell lines are among those most prominently recognized by the lectibody even at the lowest concentration (i.e., 0.1 µg/mL or 1.3 nM) analyzed **(Figure 1A)**. By contrast, AvFc poorly recognized normal human PBMCs and non-tumorigenic cell lines including MCF10 mammary gland epithelial and BEAS-2B lung epithelial cells. Marginal binding was also noted for relatively few cancer cell lines, including MDA-MB-468 breast carcinoma, Raji Burkitt’s lymphoma and SU-DHL-4 B cell lymphoma cells **(Figure 1A, B and C)**.

**Figure 1.**
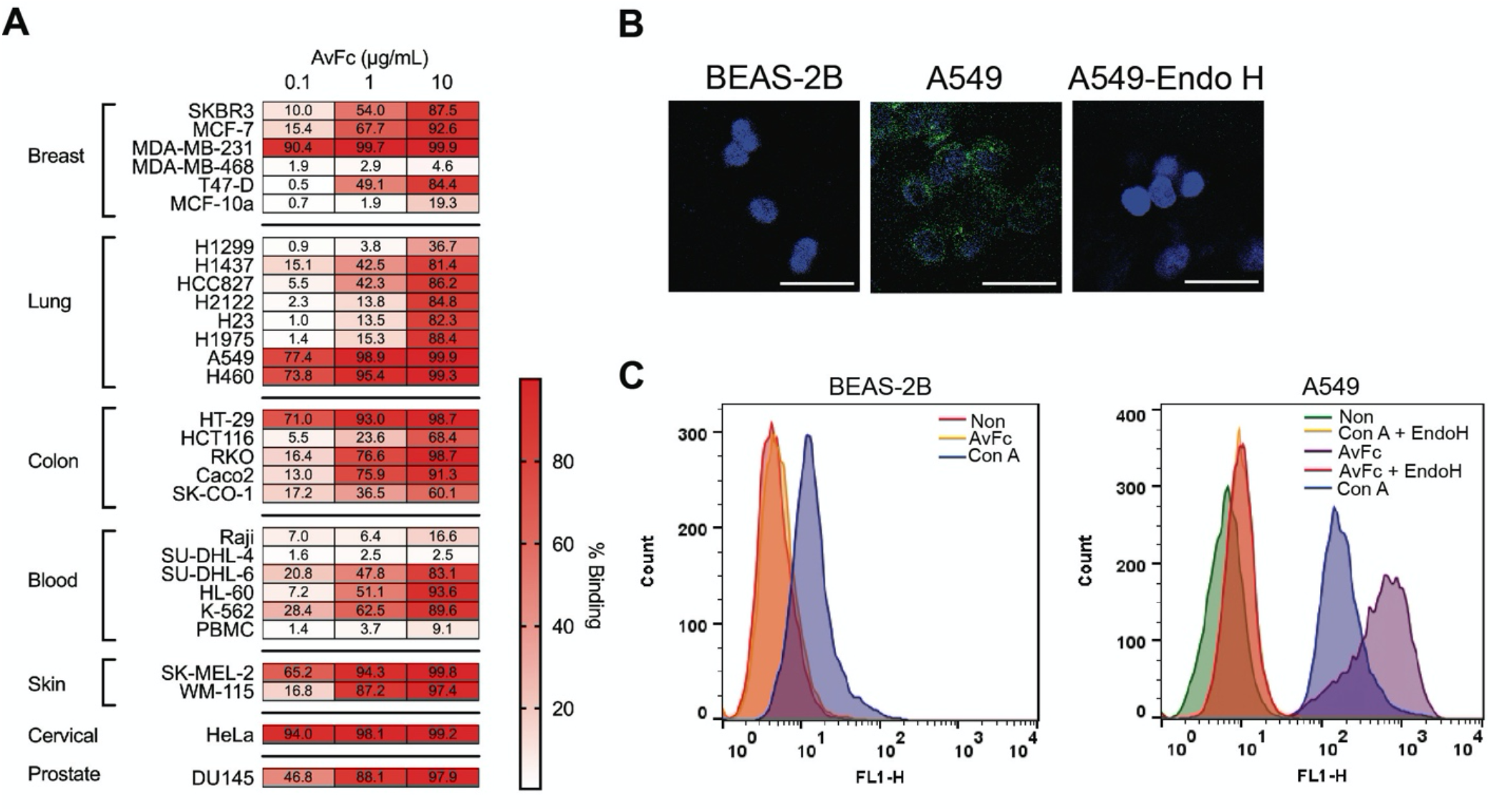
AvFc recognizes high-mannose glycans on cancer cell lines and has antiproliferative effects. (A) The binding of AvFc to various cancer cell lines was evaluated by flow cytometry with 0.1, 1, or 10 μg/mL of drug. The percentages of FITC positive cells are shown as a heatmap, with most cell lines becoming saturated at 10 μg/mL. (B) Immunofluorescence was used to visualize the binding of 1 μg/mL of AvFc to the non-tumorigenic lung epithelial cell line BEAS-2B and to A549 cells with or without endoglycosidase H (Endo H) treatments. AvFc does not show any binding to BEAS-2B or to endo H-treated A549 cells. (C) Flow cytometry of BEAS-2B and A549 after staining with either AvFc or Con A shows that Con A can weakly bind to both BEAS-2B and A549 cells, and that endo H digestion of cells abrogates binding by both lectins.

When A549 cells were treated with endoglycosidase H (Endo H), which specifically cleaves high-mannose glycans,^23^ the binding of AvFc to the cell line was almost completely abolished **(Figure 1B and C)**. Additionally, the lectibody’s binding to A549 cells was dose-dependently inhibited by yeast mannan and the HIV-1 envelope glycoprotein gp120 **(Figure S1)**. These results demonstrate that AvFc’s interaction with cancer cells is mediated via the high-mannose-binding activity of the lectibody’s lectin domain. The mannose-binding lectin concanavalin A (Con A) also strongly recognized A549 cells, and similarly to AvFc, this interaction was disrupted by Endo H digestion of cell-surface glycans. Unlike AvFc, however, Con A exhibited a relatively weak yet appreciable degree of interaction with the non-tumorigenic BEAS-2B cells (**Figure 1C**), highlighting distinct glycan recognition mechanisms between the lectibody and the canonical legume lectin.

### AvFc binds to EGFR and IGF1R and blocks their signaling

We continued our investigation using A549 and H460 cells, two representative NCSLC cells lines that exhibited high AvFc binding. To identify the molecular targets of AvFc in these cells, we employed a pull-down assay using Protein A beads conjugated with AvFc or AvFc^lec-^, the latter of which is a variant of the lectibody lacking high-mannose binding activity.^22^ Binding partners were isolated from A549 and H460 cell lysates and identified using mass spectrometry. Silver staining revealed unique proteins in the AvFc-bound fraction that were not isolated by the negative control, AvFc^lec-^ **(Figure 2A)**. Proteomics analyses of these fractions showed that AvFc recognized a large number of molecules that are found on the cell surface and in the extracellular matrix **(Table S1)**, with many of these being common between the two NSCLC cell lines. These included laminins, integrins, transporters and growth factor receptors **(Table 1)**. We focused on two major growth factor receptors, epidermal growth factor receptor (EGFR) and insulin-like growth factor 1 receptor (IGF1R), as they are known to play pivotal roles in cancer progression in NSCLC.^24-26^ To validate the proteomics results, co-immunoprecipitation immunoblot analysis was performed; the results confirmed interaction between AvFc and these receptors in both A549 and H460 cells **(Figure 2B)**. Because EGFR and IGF1R are dimerized upon ligand binding and phosphorylated to trigger pro-survival signaling cascades,^27-30^ we investigated whether AvFc can repress the onset of signal transduction by these receptors. After pre-incubation with AvFc, AvFc^lec-^ or the FDA-approved anti-EGFR monoclonal antibody cetuximab (CTX) in serum-free medium, A549 cells were treated with EGF or IGF1, and their respective receptors’ phosphorylation status were analyzed by immunoblot **(Figure 3A-D)**. The results indicated that AvFc, but not AvFc^lec-^, blocked the activation of both EGFR and IGF1R as evidenced by a decrease in band intensity of the phosphorylated forms of these receptors (pEGFR and pIGF1R). In contrast, while CTX blocked the activation of EGFR as effectively as AvFc, the EGFR-specific monoclonal antibody failed to inhibit that of IGF1R. To test whether AvFc can simultaneously block these receptors, we assessed the inhibition of major downstream signaling pathways shared between EGFR and IGF1R after treating A549 cells with a mixture of EGF and IGF1. Specifically, we evaluated activation of the AKT and MAPK pathways that are involved in cell invasion, proliferation and drug resistance.^31-34^ Immunoblot analysis showed that AvFc significantly blunted the phosphorylation of AKT and MAPK1 upon EGF and IGF1 co-treatment, whereas CTX and AvFc^lec-^ failed to show any inhibition **(Figure 3E-G)**.

**Table 1.** Shared between lung cancers (A549 and H460)

| UniProt Accession | Gene Name | Protein Name |
| --- | --- | --- |
| P00533 | EGFR | Epidermal growth factor receptor |
| Q14517 | FAT1 | Protocadherin FAT1 |
| P08069 | IGF1R | Insulin-like growth factor 1 receptor |
| P17301 | ITGA2 | Intergrin alpha-2 |
| P18084 | ITGB5 | Intergrin beta-5 |
| P26006 | ITGA3 | Intergrin alpha-3 |
| Q07954 | LRP1 | Low density lipoprotein receptor-related protein 1 |
| Q9Y666 | SLC12A7 | Solute carrier family12 member 7 |
| P56199 | ITGA1 | Intergrin alpha-1 |
| Q9NV96 | TMEM30A | Cell-cycle control protein 50A |
| Q15758 | SLC1A5 | Neutral amino acid transporter B(0) |
| Q9UNN8 | PROCR | Endothelial protein C receptor |
| P07942 | LAMB1 | Laminin subunit beta-1 |
| O15230 | LAMA5 | Laminin subunit alpha-5 |
| P55268 | LAMB2 | Laminin subunit beta-2 |
| O00468-3 | AGRN | Agrin |

**Figure 2.**
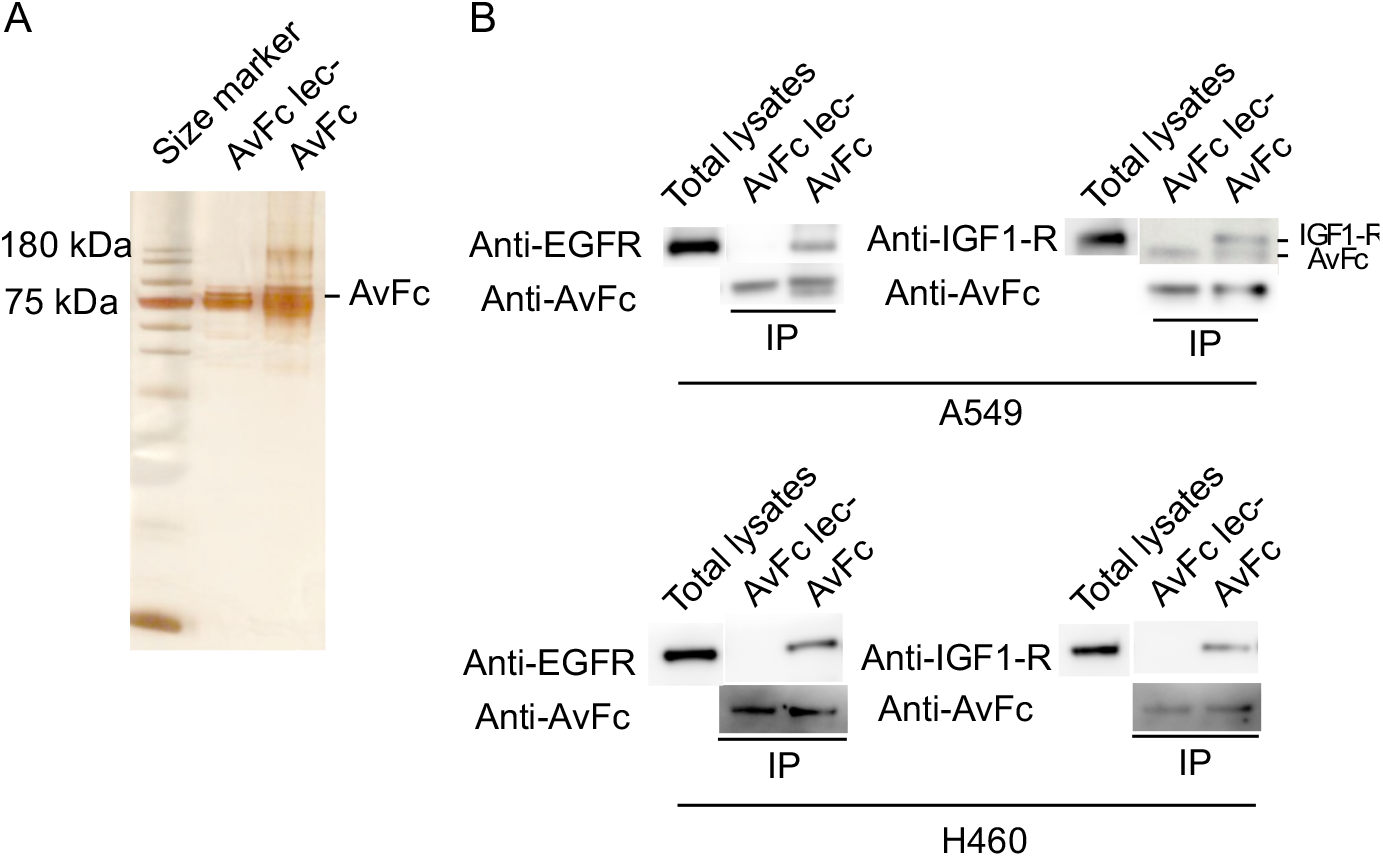
Identification of putative cancer-cell-surface binding partners of AvFc. Potential binding partners were isolated using co-immunoprecipitation and identified using mass spectrometry. (A) Silver staining of AvFc and AvFc^lec-^ fractions obtained after co-immunoprecipitation. In addition to the band corresponding to AvFc itself (≈ 77 kDa), other species at higher and lower molecular weights are present suggesting that AvFc successfully pulled down potential binding partners. (B) Co-immunoprecipitation was used to confirm the interaction between AvFc and EGFR/IGF1R isolated from A549 and H460 cells. Pull down with AvFc and then blotting with anti-EGFR or anti-IGFR antibodies revealed that AvFc, but not AvFc^lec-^, interacts with EGFR and IGF1R derived from both cell lines.

**Figure 3.**
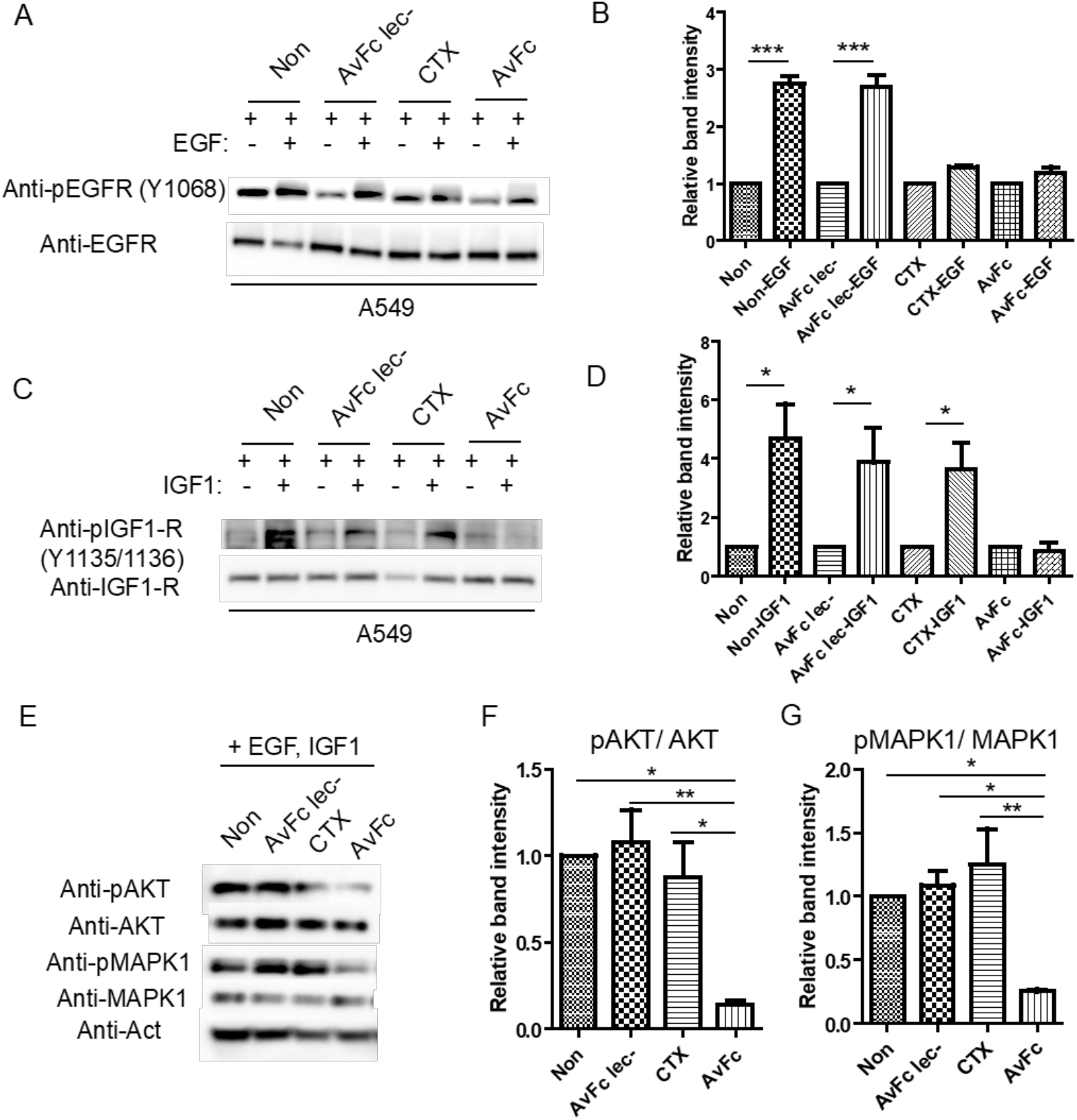
AvFc blocks EGFR and IGF1R signaling. The phosphorylation status of EGFR and IGF1R on A549 cells following treatment with their respective ligands was detected by anti-pEGFR(Y1068) and anti-IGF1R(Y1135/1136) antibodies. (A) A representative immunoblot shows that the treatment of A549 cells with 30 nM of AvFc and CTX, but not AvFc^lec-^, prior to the addition of 2 ng/ml of EGF resulted in diminished EGFR activation. (B) Quantification of immunoblot in panel A. (C) A representative immunoblot shows that only treatment of A549 cells with 30 nM of AvFc, not CTX or AvFc^lec-^, prior to the addition of 2 ng/ml of IGF1 results in decreased activation of IGF1R. (D) Quantification of immunoblot in panel C. (E) After incubation of A549 cells with AvFc, CTX or AvFc^lec-^ and subsequent stimulation with a mixuture of EGF and IGF1, AKT and MAPK1 phosphorylation was only decreased with AvFc. (F) Quantification of pAKT/AKT in panel E. (G) Quantification of pMAPK1/MAPK1 in panel E. All relative band intensities were measured by ImageJ. Bars represent mean ± SEM (N = 3). Group means were compared with the two-tailed, unpaired Student t-test for B and D, or one-way ANOVA with Bonferroni’s posttests for F and G (* *p* < 0.05, ** *p* < 0.01, *** *p* < 0.001).

### AvFc inhibits migration of A549 and H460 cells

The above results indicate that AvFc effectively binds to both EGFR and IGF1R on the cell surface, thereby intercepting their ligands and preventing receptor activation and subsequent AKT and MAPK signaling. Because AKT and MAPK signaling pathways are involved in migration,^35, 36^ we investigated the effects of AvFc on cell migration using A549 and H460 cells using a transwell culture system, wherein cells were co-incubated with the lectibody and subsequently treated with EGF or IGF1 in serum-free medium. These cells were then seeded into transwells and after 6 h cells in the bottom chamber were quantified. As shown in **Figure 4**, AvFc significantly blocked the migration of both cell lines regardless of which growth factor the cells were stimulated with, whereas AvFc^lec-^ failed to show any effect. CTX, on the other hand, only effectively inhibited the migration of A549 cells treated with EGF, but not when the cells were treated with IGF1 **(Figure 4A)**. Additionally, CTX failed to show any effect on the migration of H460 cells, even when the cells were stimulated with EGF **(Figure 4B)**.

**Figure 4.**
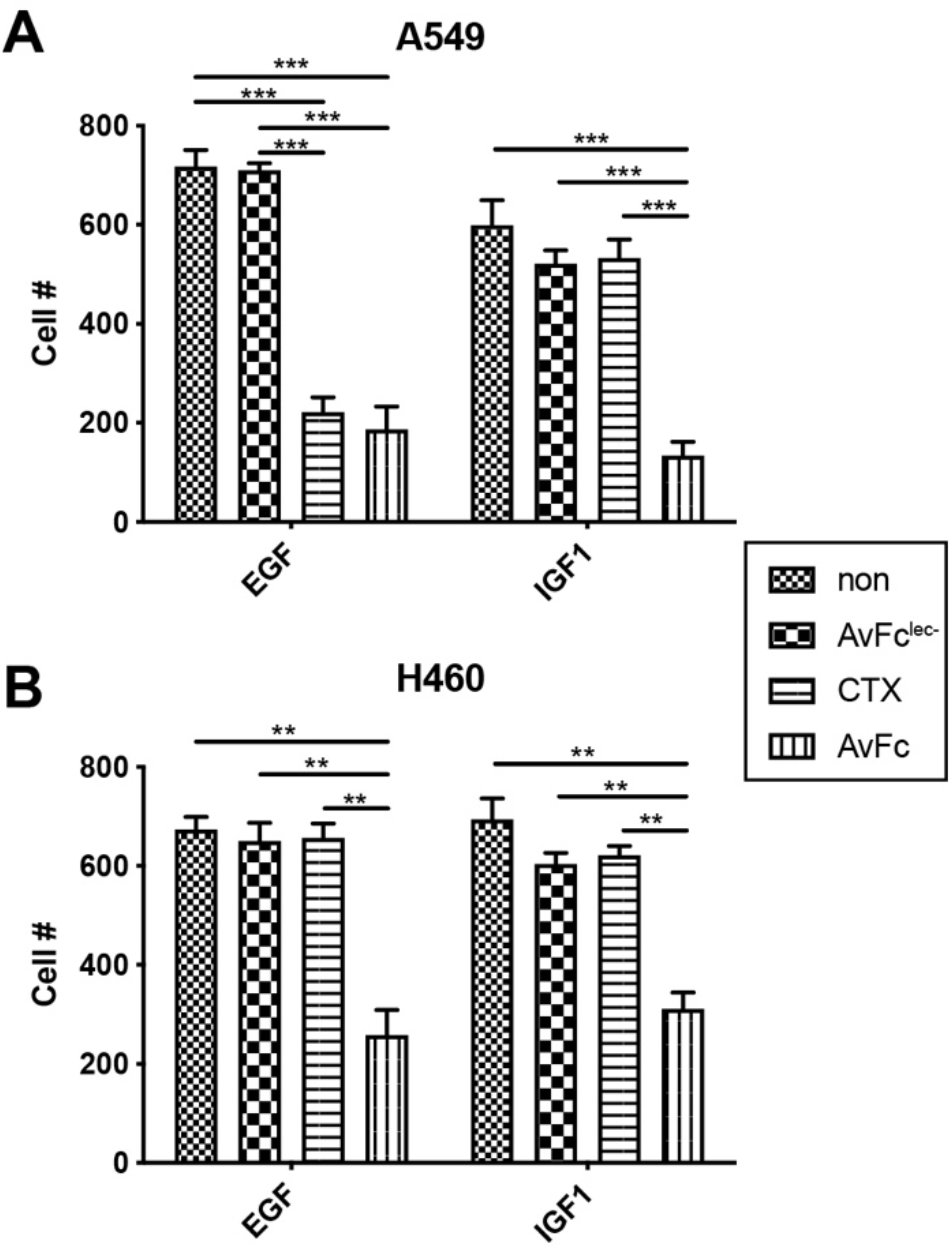
AvFc inhibits A549 and H460 cell migration. Migration of (A) A549 cells and (B) H460 cells was measured in transwells with 8 μm pores after treatment of cells with 30 nM of AvFc, CTX, or AvFc^lec-^ and stimulation with EGF or IGF1. Error bars represent mean ± SEM from three replicates. Groups were analyzed by two-way ANOVA followed by Bonferroni’s Multiple Comparison Test (** *p* < 0.01, *** *p* < 0.001).

### AvFc induces ADCC against cancer cells

Next, we investigated the consequences of AvFc’s binding to cancer cells from a different angle, asking whether the Fc region of the lectibody can direct antibody-dependent cell-mediated cytotoxicity (ADCC). Since ADCC is mediated primarily through NK cells expressing FcγRIIIa, we assessed the Fc-mediated activity of AvFc against A549 and H460 cells using an *in vitro* assay based on FcγRIIIa-activated luciferase expression. As shown in **Figure 5A**, AvFc activated FcγRIIIa in a dose dependent manner. AvFc^lec-^, on the other hand, failed to activate FcγRIIIa at all concentrations tested, demonstrating that the activity is dependent upon AvFc’s binding to high-mannose glycans on cancer cells. Of note, AvFc showed significantly higher efficacy against A549 cells than CTX at the top three concentrations tested (0.08, 0.40 and 2.00 µM). Moreover, AvFc exhibited remarkable activity (a maximum over 30-fold above baseline) against H460, while CTX was ineffective for the large-cell lung carcinoma cell-line **(Figure 5A)**. To confirm the ability of AvFc to induce Fc-mediated cell killing activity, a canonical ADCC assay was performed using human PBMC effector cells and A549 cells as the targets. As shown in **Figure 5B**, the lectibody showed a dose-dependent cell lysis activity against A549 cells; it is of note, however, that the efficacy of the lectibody was nearly twice as high as that of CTX (∼80% for AvFc vs. ∼40% for CTX).

**Figure 5.**
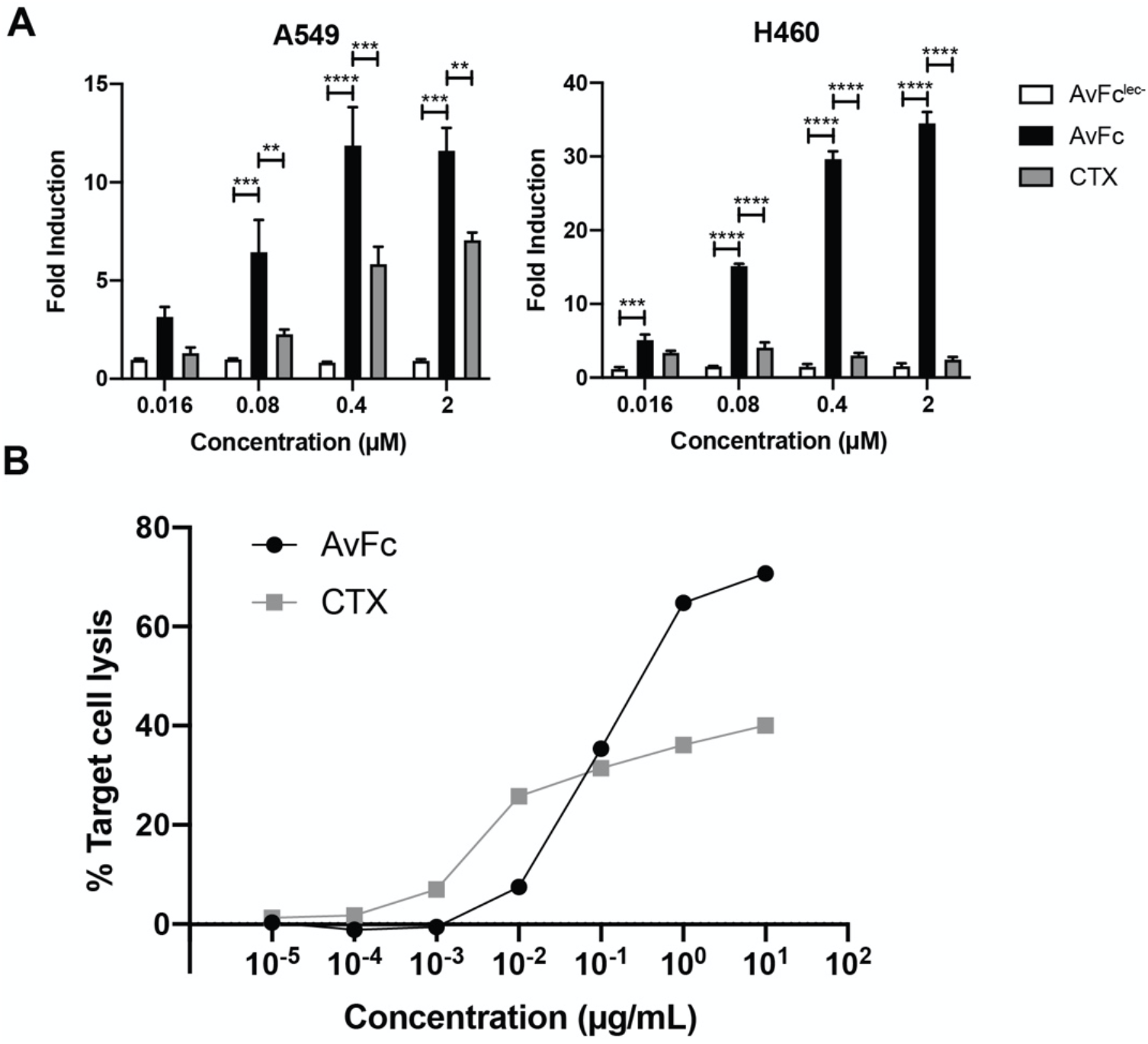
*In vitro* Fc-mediated anticancer activity of AvFc. (A) FcγRIIIa activation by AvFc, AvFc^lec-^ and CTX against A549 or H460 cells in a reporter cell-based assay. Representative data from at least two independent experiments are shown. Columns and bars represent mean ± SEM (n=3). Groups were compared with two-way ANOVA followed by Tukey multiple comparison test (** *p* < 0.01, *** *p* < 0.001, **** *p* < 0.0001). (B) A human PBMC-based ADCC assay using A549 lung cancer cells. Cells were pre-incubated with serial dilutions of AvFc or CTX for 30 min in a 37°C/5% CO2 incubator. PBMCs were added to initiate the ADCC effects at an optimized effector/target ratio (50:1 for AvFc, 25:1 for Erbitux). After incubation in a 37°C/5% CO2 incubator for 6 h, cell supernatants were collected for measuring released lactose dehydrogenase to calculate % target cell lysis. The experiment was done in triplicates, with mean ± SEM shown for each data point.

### AvFc exhibits antitumor effects in mouse A549 and H460 xenograft models

The anti-tumor effects of AvFc were evaluated in Prkdc^scid^/SzJ (SCID) mice challenged with A549 and H460 xenografts implanted in the hind left flank. Intraperitoneal treatment with 25 mg/kg of AvFc or CTX was initiated at day 4 post tumor challenge and continued every two days for a total of 6 doses. AvFc treatment significantly blunted A549 **(Figure 6A)** and H460 **(Figure 6B)** tumor growth compared to the vehicle control. On the other hand, CTX showed similar efficacy to AvFc against A549 tumors but failed to show an effect on the growth of H460 tumors. To further evaluate AvFc’s anti-tumor effect, mice were intravenously challenged with A549-GFP and subsequently treated with the same dosing regimen as in the flank tumor models. Fluoresence imaging of the lung isolated 18 days after the last dose showed that AvFc significantly inhibited the growth of A549-GFP cells in the lung as compared to a vehicle control **(Figure 6C)**. Taken together, these data clearly demonstrated that AvFc has the ability to elicit antitumor activity *in vivo*.

**Figure 6.**
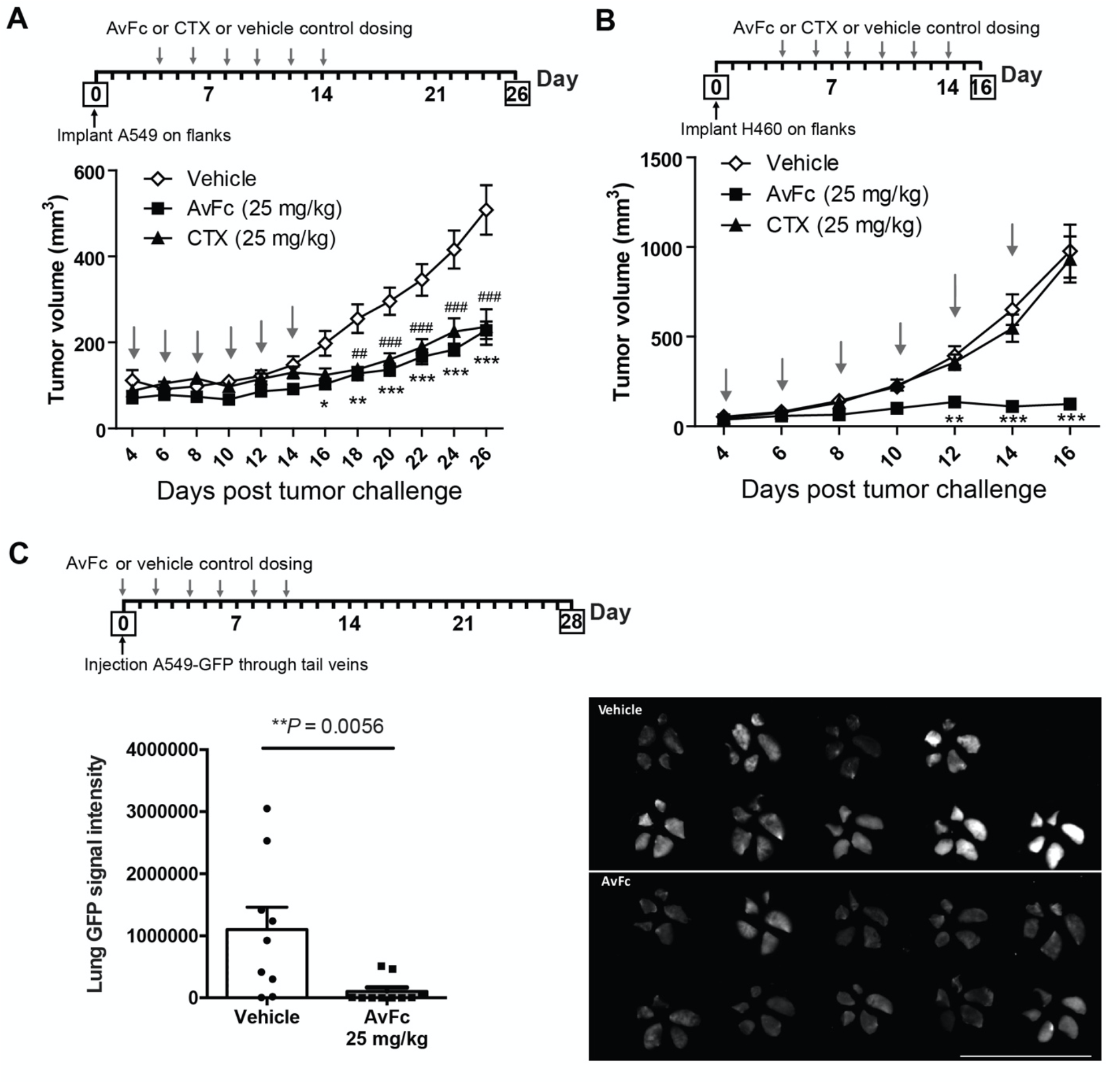
*In vivo* anticancer activity of AvFc. (A) The A549 subcutaneous xenograft challenge model and (B) the H460 subcutaneous xenograft challenge model in SCID mice. Four days post challenge, mice were treated i.p. with AvFc or CTX at 25 mg/kg, or a vechicle control every other days (Q2D) for a total of 6 doses, as indicated by arrows. Animals were monitored until day 26 for A549 and day 16 for H460 models. Tumor volumes were compared with two-way ANOVA followed by Tukey multiple comparison tests (* *p* < 0.05, ** *p* < 0.01, *** *p* < 0.001 between vehicle and AvFc; # *p* < 0.05, ## *p* < 0.01; ### *p* < 0.001 between vehicle and CTX). (C) The A549-GFP human lung cancer metastasis model in SCID mice. SCID mice were i.v. challenged with A549-GFP cells on day 0, followed by every-other-day dosing with an i.p. injection of AvFc at 25 mg/kg (n=10) or a vehicle control (n=9), as indicated in the diagram. On day 35, the lungs were removed and GFP signal intensity of the lung from each mouse was quantified. Columns and bars represent mean ± SEM, with dots representing individual mice. GFP images of the lungs (four right and single left lobes) from all animals in vehicle and AvFc groups are shown in the right. Bar = 5 cm. Statistical difference between groups was analyzed by the Mann-Whitney test.

### AvFc preferentially binds to human NSCLC tumor tissue

Lastly, we investigated the binding of AvFc to primary tumor and adjacent tissues isolated from human NSCLC patients using immunohistochemistry (IHC). Compared to the adjacent tissue, AvFc binding was more evident in NSCLC tumor (**Figure 7A**), indicating that the lectibody is capable of distinguishing their differential glycosylation patterns. Among the matched pair tissues from 10 NSCLC patients analyzed, 7 showed significantly higher AvFc binding in tumors than in the adjacent tissue **(Figure 7B)**. Given that EGFR was one of the major molecular targets of AvFc in A549 and H460 (Figure 2), we postulated that AvFc’s tumor selectivity found in NSCLC patients’ lung tissues may be partly attributed to the receptor. Thus, EGFR was enriched from the tumor and adjacent tissue lysates from five NSCLC patients (sample No. 151, 117, 448, 234, 155) using co-immunoprecipitation and then detected with AvFc, CTX, or AvFc^lec-^ by Western blot. A representative image of tissue samples from Patient 117 is shown in **Figure 7C**, and relative band intensities between tumor EGFR and the adjacent tissue-derived counterpart are shown in **Figure 7D**. Although AvFc reacted with both tumor and adjacent tissue EGFRs, it showed a stronger signal with the former (approximately 2-fold on average; **Figure 7D**), indicating that the lectibody had higher affinity to tumor-derived EGFR. By contrast, CTX showed bands of similar intensity for tumor and adjacent tissue EGFRs in all five tissues tested (**Figure 7C,D**). AvFc^lec-^ failed to probe the receptor from both tumor and adjacent tissues. Taken together, these results indicate that AvFc has preferential binding to NSCLC tumor-derived EGFR over that of the normal lung tissue, while CTX cannot distinguish them.

**Figure 7.**
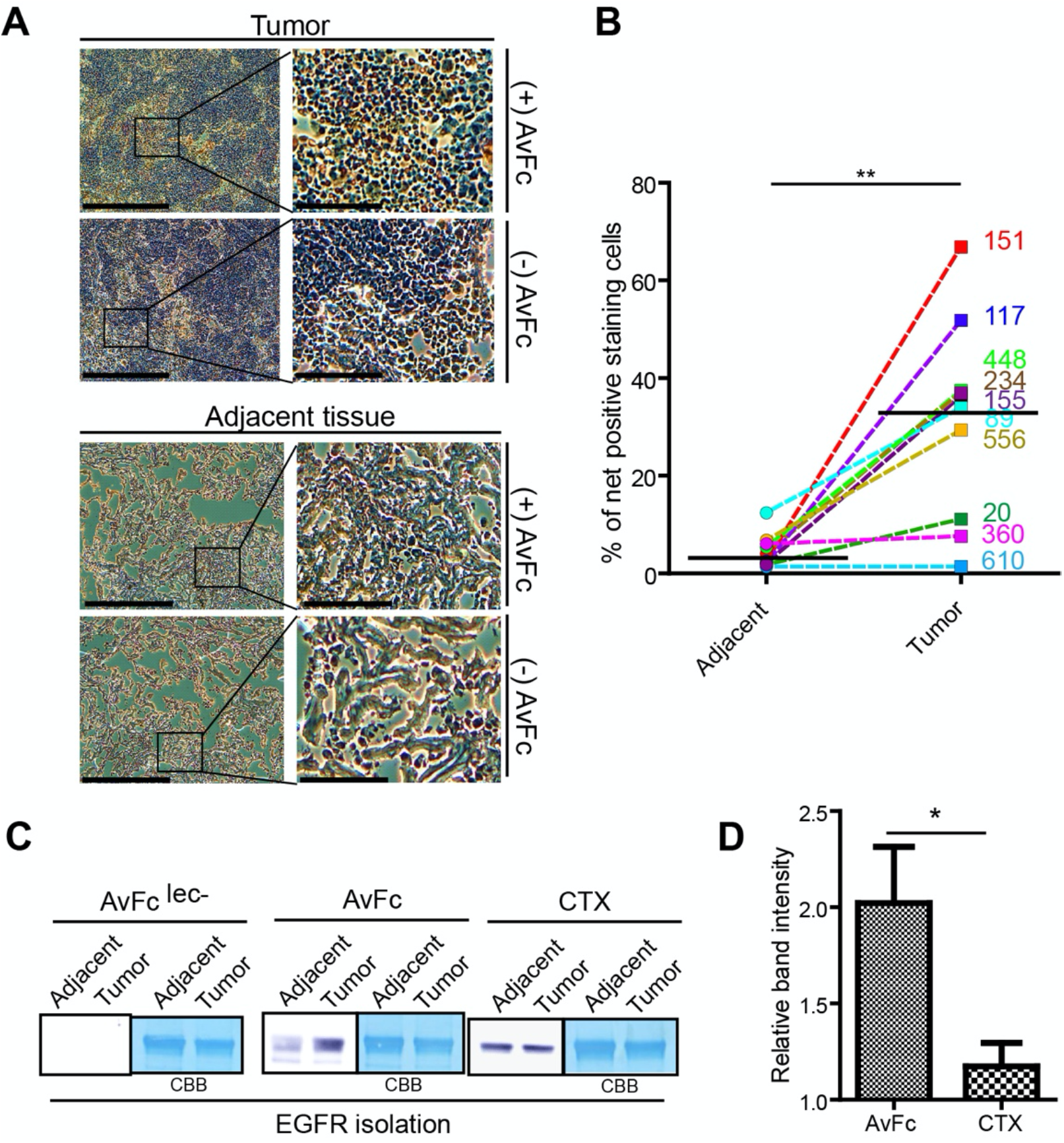
Analysis of AvFc binding to primary human lung tissue and EGFR. Binding to human NSCLC tissue was evaluated using IHC. (A) IHC staining with AvFc or a biotinylated anti-human IgG secondary antibody only. Representative stains from Patient 117 lung tissues are shown, with hematoxylin as a counter stain. Bar = 200 µm (left image) or 100 µm (right image). (B) Quantification of AvFc staining for lung tissues from all 10 patients tested using ImageJ. The number of positively stained cells between tumor and matched adjacent tissue was compared using the non-parametric Wilcoxon matched-pairs signed rank test (** *p* < 0.01). (C) Representative immunoblot analysis of EGFR isolated from NSCLC tumor or matched adjacent tissue samples from 5 patients. EGFR was isolated by anti-EGFR IgG1 with Protein A bead precipitation and detected with AvFc^lec-^, AvFc or CTX. (D) Quantification of EGFR immunoblot using a densitometory analysis. Relative binding intensities (tumor:adjacent) are shown for AvFc and CTX. Columns and bars represent mean ± SEM (n=5), and data were compared using an unpaired t-test (\**p* < 0.05).

## DISCUSSION

Aberrant glycosylation has long been recognized as a hallmark of cancer. Nevertheless, development of therapeutics targeting cancer-associated glycans has been slow. In the present study, we showed that AvFc, a lectibody specific to high-mannose glycans,^19^ can recognize multiple human cancer cell lines derived from various cancer types. The therapeutic implications of AvFc’s interaction with cancer cells were evaluated using two NSCLC cell lines A549 and H460, demonstrating that the lectibody can block the activation of EGFR and IGF1R and cell migration upon stimulation with their respective ligands, elicit ADCC activity, and significantly delay xenograft tumor growth in SCID mice. Furthermore, IHC analysis showed that AvFc preferentially binds to primary human NSCLC tumors over adjacent non-tumor lung tissues isolated. To our knowledge, this is ths first report demonstrating the antitumor effects of an agent targeting cancer-associated high-mannose glycans.

An initial analysis indicated that AvFc has high selectivity to malignant cells over noncancerous or normal healthy cells, since AvFc did not show any significant binding to nontumorigenic epithelial cell lines MCF10a and BEAS-2B, human PBMCs **(Figure 1A-C)**, or primary mesenteric lymph node cells isolated from rhesus macaques.^19^ Similar results regarding the selectivity for cancer-associated high-mannose glycans were previously reported with TM10, an IgM monoclonal antibody isolated from mice immunized with FasL-expressing B16F10 mouse melanoma cells.^37^ Similar to AvFc, the epitope of TM10 appeared to be clusters of high-mannose glycans, in particular Man9, and the antibody recognized human melanoma, prostate, ovarian, and breast cancer cells with no apparent surface binding to untransformed cells. However, unlike AvFc, TM10 showed little *in vivo* or *in vitro* anticancer activity. The authors attributed the lack of therapeutic effects to the specific isotype of TM10 antibody given that antibodies of IgM isotype typically have poor tissue penetration, short biological half-lives, and lack Fc-mediated effector functions.^37^ Thus, while it appears that selectivity for cancer cells is similar between AvFc and TM10, the presence of the Fc region from IgG1 is the major differentiating factor of AvFc, a molecular design that offers significant advantages as a potential anti-cancer agent.

Our findings in the present study add to growing evidence indicating that a high proportion of high-mannose glycans represents a unique characteristic of the cancer cell glycocalyx. Unlike other conventional mannose-binding lectins like Con A, AvFc preferentially recognizes clusters or groupings of high-mannose glycans containing terminal α1,2-linked mannose residues.^19, 22^ Such a high density of high-mannose structures is rare in the glycocalyx of normal cells, as illustrated by our data showing AvFc’s inability to recognize the nontumorigenic BEAS-2B bronchial epithelial cell line while Con A showed noticeable binding (**Figure 1C**). Con A interacts with both internal and external α-D-mannosyl and α-D-glucosyl residues and has four sugar-binding sites.^38^ As such, it is more “promiscuous” than AvFc and capable of recognizing a broader spectrum of glycoforms. Furthermore, preliminary data suggest that AvFc has low affinity to individual glycans and glycoproteins with small numbers of glycans but high affinity to high-mannose rich glycoproteins like HIV gp120 (Dent et al. manuscript in prepration). This implies that there exists a threshold level of high-mannose glycans that must be present in proximity in order for AvFc to bind with any appreciably affinity, and non-cancer cells simply may not reach this threshold. Although such a threshold level is unclear at the moment and needs further invenstigation, the data herein support the notion that AvFc is superior to conventional mannose-specific lectins with respect to selectivity to tumor-associated high-mannose glycans.

Given that changes in *N*-glycosylation modification would globally give rise to any glycoproteins in cancer cells, it is not surprising that the proteomics analysis revealed a large number of proteins recognized by AvFc in A549 and H460 cells **(Table 1)**. Of interest is that EGFR and IGF1R were among the common cell-surface glycoproteins targeted by AvFc in these NSCLC cells **(Figure 2)**. The result indicates that these receptors, which are often overexpressed and strongly associated with cancer progression in NSCLC,^24-26^ display dense high-mannose glycans on the cancer cells. In fact, they are both highly glycosylated, containing as many as 13 and 16 *N*-glycosylation sites (UniProtKB: P00533 and P08069), respectively. Also, since the ectodomains of EGFR in its activated form and IGF1R exist as dimer,^39-41^ it is plausibile that AvFc exhibits high affinity for these receptors. In **Figure S2**, we found that AvFc has higher affinity to EGFR dimer than to EGFR monomer, which suggests that high expression and activation of EGFR on some cancer cell surfaces may increase local high-mannose glycan concentrations and facilitate AvFc’s binding and antitumor activity. One of the consequenes of AvFc binding to EGFR and IGF1R was the blockade of their activation and subsequent downstream signaling, as demonstrated by the data showing that AvFc inhibited the phosphorylation of both receptors as well as AKT and MAPK1 upon stimulation with their respective ligands in A549 cells (**Fig. 3A-G**).

AvFc’s ability to inhibit both EGFR and IGF1R has important therapeutic implications, as currently there is no FDA-approved anticancer drug that can simultaneously block these receptors. CTX, a FDA-approved anti-EGFR antibody therapeutic used in the present study as a reference, was only able to block EGFR but not IGF1R in A549 (**Fig. 3, 4**) and, in stark contrast to AvFc, could not exhibit any antitumor effect against the H460 cell line (**Fig. 4-6**), which is known to be CTX resistant.^42^ Despite the fact that colorectal cancer patients show significant improvements in response rates, overall survival and progression-free survival after CTX treatment, CTX responses in NSCLC patients have not been convincing in clinical trials.^43-45^ IGF1R may be involved in the resistance mechanism of CTX and other EGFR-targeted drugs,^46- 49^ since these two receptors share similar downstream signaling pathways (PI3K/AKT/MAPK/NF-κB); IGF1R can bypass EGFR inhibition, while their cooperation may promote tumor growth and progression.^48-50^ One study revealed that overexpression of both EGFR and IGF1R was observed in 24.8% of 125 surgical NSCLC patients, and high co-expression of EGFR and IGF1R was a significant prognostic factor of worse disease-free survival.^50^

To further elucidate the potential anti-tumor mechanism of action of AvFc, we measured ADCC activities in both reporter cell-based assays and human PBMC-based assays (Figure 5A and B). In both assay formats AvFc elicited a strong ADCC response by effectively activating FcγRIIIa on the surface of engineered Jurkat cells (which express luciferase in response to activation) and PBMCs. Conversely, CTX had no activity against H460 cells and only moderate activity against A549 in both the reporter cell assay and PBMC-based assay, underperforming AvFc. This was consistent with the anti-tumor effects of AvFc seen in Prkdc^scid^/SzJ (SCID) mice challenged with A549 and H460 xenografts (Figure 6A-C), wherein AvFc treatment slowed the growth of both xenografts while CTX was only efficacious against A549. Taken together, these results suggest that binding and inhibition of multiple cell-surface receptors in addition to more potent ADCC activity resulted in AvFc’s superior anticancer activity in these models.

The selectivity of AvFc was evaluated in 10 pairs of tumor and adjacent lung tissues from NSCLC patients (Figure 7A-D). Overall, AvFc interacted preferentially with tumor tissue and was capable of distinguishing the tumor and adjacent tissues, despite the low level of background AvFc staining in the latter. In this initial study we did not investigate high-mannose expression levels within the tissues, which will likely depend on the developmental stage of the cancer, nor did we evaluate EGFR gene mutations. Despite these limitiations, the selective interaction of AvFc with primary NSCLC cells in this analysis demonstrates the utility of AvFc beyond animal models and suggests that it may be able to effectively target tumors in NSCLC patients.

While our findings lend support for the notion that high-mannose glycans constitute a cancer glycobiomarker, it remains elusive how or why high-mannose glycans become over-represented on the surface of cancer cells in the first place. A few cellular mechanisms have been identified which allow the accumulation of immature glycans in tumors, including stress-independent activation of X-box binding protein 1 (XBP1), misregulation of N-acetylglucosaminyltransferase, and downregulation of α-mannosidase I and mannosyl(α-1,3-)-glycoprotein β-1,2-N-acetylglucosaminyltransferase (MGAT1).^11, 12, 51, 52^ Stress-independent activation of XBP1, for instance, was found to reduce sialylation and bisecting GlcNAc while increasing the levels of high-mannose glycans in HEK293 and HeLa cells by affecting mannosidase expression.^51^ The significant reduction of α-mannosidase I expression found in cholangiocarcinoma cells was correlated to the elevation of high-mannose glycans and a more metastatic phenotype.^12^ The use of kifunensine, a small-molecule inhibitor of α-mannosidase I, also increased high-mannose glycan content and produced similar results.^12^ Takayama et al. have shown that increased high-mannose glycan expression, detected in surgical specimens of hepatocellular carcinoma, was associated with decreased expression of MGAT1, a key glycosyltransferase that converts high-mannose glycans to complex- or hybrid-type N-glycans.^11^ Meanwhile, high-mannose glycans at the helical domain of transferrin receptor protein 1 appear to trigger structural changes that improve noncovalent interaction energies, resulting in cell migration enhancement in metastatic cholangiocarcinoma.^12^ A recent publication assessing the impact of high-mannose glycans on bone-marrow-derived mesenchymal stromal cells has provided evidence that these glycans alter the physical and structural properties of the cells themselves, decreasing their size and increasing motility, which may in part explain the greater metastatic potential seen in other cell lines.^53^ Given the growing body of evidence indicating the close association between the abundance of high-mannose glycans on cancer cells and increased cell migration and metastatic potential, it is of high clinical significance to uncover the cause and process leading to high-mannose overexpression in cancer and to scrutinize its functions in tumor microenvironments and metastasis. In this regard, AvFc may provide a valuable tool to probe and monitor high-mannose glycan accumulation on cell surface, thereby facilitating such investigations.

In summary, the present study demonstrated that AvFc, a lectibody targeting high-mannose glycans, can selectively recognize cancer cells and exert antitumor activity possibly through a combination of growth factor receptor inhibition and immune activation via Fc receptors. Our findings suggest that increased abundance of high-mannose glycans in cancer, such as those expressed on EGFR and IGF1R, can be a druggable target in NSCLC, and AvFc may provide a new tool to probe and target tumor-associated high-mannose glycan biomarker.

## MATERIALS AND METHODS

### Human lung tissues

We acquired de-identified post-operative human lung cancer tissues and paired adjacent tissues from University of Louisville hospital (Louisville, KY). The pathological type of each tumor was determined to be NSCLC. Informed written consent was provided by all participants and the study protocol was approved by the Human Subjects Protection Program of University of Louisville (Study #18.1240). Tissues were immediately frozen in liquid nitrogen at the surgery and stored at −80°C.

### Animal housing and care

Nine-week-old female Prkdc^scid^/SzJ (SCID) mice (The Jackson Laboratory, Bar Harbor, ME) were housed in a temperature-controlled environment, with an alternating light/dark cycle of 12 hours and free access to standard diet and water. The investigators were not blinded for sample administration. All experimental procedures were approved by the University of Louisville’s Institutional Animal Care and Use Committee.

### Reagents

Antibodies specific to EGFR (D38B1), phospho-EGFR (Y1068), IGF1R (D23H3), phospho-IGF1R (Y1131), AKT, phospho-AKT (S473), MAPK1, and phospho-MAPK1 (ERK1/2) were purchased from Cell Signaling Technology (Danver, MA). EGF and IGF1 were purchased from Thermo Fisher Scientific (Waltham, MA). CTX was obtained from the University of Louisville Hospital pharmacy.

### Cell culture

All cell lines were obtained from American Type Culture Collection (ATCC, Manassas, VA) and authenticated by the supplier. Cells were grown according to ATCC’s recommendations, regularly screened for mycoplasma using a commercial PCR-based kit (ATCC) and tested at low passage numbers, with quality ensured based on viability and morphologic inspection. In particular, A549 cells were grown in DMEM supplemented with 10% FBS and 1% penicillin/streptomycin and H460 cells were grown in RPMI 1640 supplemented with 10% FBS and 1% penicillin/streptomycin unless otherwise stated.

### Production of AvFc

AvFc and AvFc^lec-^ were produced using a transient plant expression vector in *Nicotiana benthamiana* as described previously.^19^ Briefly, 4 week old plants were transformed with a magnICON® vector containing the gene for AvFc by agroinfiltration and incubated for one week. At that time, leaf tissue was homogenized in a NaPi buffer at a pH of 7.4 and clarified by centrifugation, followed by fast protein liquid chromatography on the ÄKTA pure system (GE Healthcare Life Sciences, Chicago, IL) using protein A as the first chromatography step and ceramic hydroxyapatite (CHT) as a cleanup step. Endotoxin was removed from the purified protein using the Triton X-114 phase separation method, followed by concentration of the protein using a 10 kDa MWCO centrifuge filter and sterilization with a 0.2 μm filter. Purity was assessed with SDS-PAGE, with AvFc appearing as a band at approximately 77 kDa under non-reducing conditions.

### Flow cytometry analysis of AvFc binding to human cells

Cancer-cell lines were harvested and incubated with various concentrations of AvFc (0.1, 1 and 10 µg/mL) in culture medium for 30 minutes on ice and washed 3 times with DPBS. Cells were then incubated with goat F(ab’)2 anti-Human IgG Fc-FITC antibody (Abcam, Cambridge, MA) for 30 minutes in the dark on ice. After washing 3 additional times with DPBS, the cells were fixed with 1% formalin for 15 minutes on ice. Data were acquired on a FACSCalibur flow cytometer (BD BioSciences, San Jose, CA) by counting 10,000 events per sample and determining the percentage of FITC^+^ cells with FlowJo. The non-sugar-binding mutant AvFc^lec-^ was used as a negative control. The analyses were performed in triplicate.

### Immunofluorescence

1,000 A549 cells or 10,000 BEAS-2B cells were seeded per chamber in Lab-Tek II chamber slides (Thermo Fisher Scientific, Waltham, MA) and incubated for 24 hours. After washing with PBS, cells were fixed in 4 % formalin in PBS for 20 minutes at room temperature. After incubation with 0.2% Triton X-100 in PBS for 15 minutes at room temperature, Human Fc Block™ (BD, San Jose, CA) was added to cells and incubated for an additional 10 minutes. Cells were then blocked with 3% BSA-PBS for 30 minutes at room temperature and then incubated with 250 units of endoglycosidase H at 37°C for 1 hour, according to the manufacturer’s protocol (New England Biolabs, Ipswich, MA). Cells were then stained with 10 µg/ml of AvFc for 3 hours at room temperature and, after washing with PBS, stained with a 1:40 dilution of anti-human IgG-FITC (Sigma, Mendota Heights, MN) for 1 hour at room temperature. Cells were then mounted with coverslips using mounting medium for fluorescence with DAPI (VECTASHIELD®, Burlingame, CA). Slides were analyzed by fluorescent confocal microscopy (ZEISS LSM 880).

### Co-immunoprecipitation

1×10^6^ A549 or H460 cells were seeded in a 10 cm^2^ plate (Corning, Tewksbury, MA) and incubated in growth medium for 24 hours. Cells were washed with PBS and cell lysates were prepared in T-PER buffer (Thermo Fisher Scientific, Waltham, MA) supplemented with a protease/phosphatase inhibitor (Thermo Fisher Scientific, Waltham, MA). After centrifugation at 13,000 xg for 10 minutes at 4°C, supernatants were mixed with 4 µg of AvFc or AvFc^-lec^. After incubation for 24 hours at 4°C, 10 µl of protein A beads (Santa Cruz, Dallas, TX) were added. After an additional incubation for 2 hours at 4°C, the mixture was washed with T-PER buffer and immune-blot analysis was performed.

### EGFR isolation

Tissue homogenates were prepared by silicon beads and Precellys® 24 homogenizer (Bertin, Rockville, MA) in T-PER buffer (Thermo Fisher Scientific, Waltham, MA) with protease inhibitor cocktail (Sigma, Mendota Heights, MN). Debris were removed by centrifugation at 13,000 xg for 10 minutes at 4°C. Supernatants were incubated with 4 µg of Anti-EGFR IgG1 (D38B1) (Cell Signaling Technology, Danver, MA) and 20 µg of protein A beads (Santa Cruz, Dallas, TX) for 4 hours at 4°C. The mixture was washed with T-PER buffer.

### Immunoblot analysis

SDS-PAGE and membrane transfer cassettes were purchased from Thermo Fisher Scientific (Waltham, MA). Protein samples were run on 10% Bolt Bis-Tris Plus gels with NuPAGE MES SDS running buffer (Thermo Fisher Scientific, Waltham, MA). Transfer to PVDF membranes in NuPAGE transfer buffer was carried out at 10 V overnight at 4°C. Membranes were then incubated in 3% BSA in TBST for 2 hours and anti-EGFR, anti-IGF1R, anti-human Fc, or AvFc in TBST supplemented with 1% BSA were incubated over-night at 4°C. HRP tagged secondary antibodies (Anti-rabbit IgG, Santa Cruz, Dallas, TX, Anti-human IgG, SouthernBiotech, Birmingham, AL) were used for the protein detection and membrane images were taken using Amersham Imager 600 (GE Healthcare Life Sciences, Chicago, IL).

### Proteomics analysis

Potential cell-surface binding partners of AvFc were identified using co-immunoprecipitation followed by mass spectrometry. Co-immunoprecipitation in this instance was performed using the Pierce Co-immunoprecipitation Kit (Thermo Fisher Scientific, Waltham, MA) according to the manufacturer’s instructions. Briefly, whole-cell lysates of A549 and H460 cells were pre-cleared with a control agarose resin to reduce non-specific interactions, then co-incubated with 100 μg of AvFc or AvFc^lec-^ which were covalently attached to an agarose resin for 2 hours at 4°C. Bound proteins were then eluted using a low pH buffer and neutralized with 1 M tris base for mass spec analysis.

Protein samples were digested with trypsin (1:50 ratio) in a filter-aided sample preparation approach following reduction and alkylation with 100mM dithiothreitol and 50mM iodoacetamide. The tryptic digests (0.5µg) were separated using a Proxeon EASY n-LC (Thermo-Fisher Scientific) UHPLC system and Dionex (Sunnyvale, CA) 2cm Acclaim PepMap 100 trap and a15cm Dionex Acclaim PepMap RSLC (C18, 2µm, 100Å) separating column. The eluate was introduced into an LTQ-Orbitrap ELITE (Thermo-Fisher Scientific) using a Nanospray Flex source and MS2 data collected in a data dependent fashion in a top-20 rapid CID method. All MS1 data were acquired using Fourier transform ion cyclotron resonance MS at 240,000 resolution and MS2 data using the linear ion trap. MSn data were searched using Proteome Discoverer 1.4 (Thermo Scientific) with Sequest HT (SageN) and Mascot, version 4.0 (Matrix Science) in a decoy database search strategy against UniProt Knowledgbase, Homo sapiens reference proteome. The searches were performed with a fragment ion mass tolerance of 1.0 Da and a parent ion tolerance of 50 ppm. The search data results file were imported into Scaffold, version 4.3.4 (Proteome Software Inc.) and filtered using a 2ppm mass error filter, removal of decoy hits, to control for <1.0% false discovery rates with PeptideProphet and ProteinProphet (Institute for Systems Biology). Peptide and protein identifications were accepted at >95.0% probability by the PeptideProphet or ProteinProphet algorithm. A comparison of protein abundance among the sample sets was conducted in Scaffold using the intensity based absolute quantification (iBAQ) method. Results were further refined using Gene Ontology (GO) terms to extract the most abundant membrane receptors, transporters, and adhesion molecules bound by AvFc and not AvFc^lec-^.

### Transwell migration assay

1×10^5^ A549 or H460 cells in 200 µl of serum-free growth medium were seeded in the insert of a transwell plate with 8 μm pores (VWR International, Radnor, PA). These cells were coincubated with AvFc or CTX at 30 nM for 2 hours at 37 °C. Afterwards, growth medium supplemented with 20% FBS was added to the outside well and EGF or IGF-1 was added to a final concentration of 2 ng/mL in the transwell insert. After 6 hours, migrated cell counts were determined by trypsinization and trypan blue staining (Thermo Fisher Scientific, Waltham, MA).

### ADCC Reporter Assay

Antibody-dependent cell-mediated cytotoxicity (ADCC) was assessed by an ADCC Reporter Bioassay (Promega, Madison, WI) following the manufacturer’s protocol. Each sample was tested in triplicate. Briefly, 3 NSCLC cell lines used as target cells were seeded in an opaque white 96-well flat-bottom culture plate (Corning, Tewksbury, MA) at 10,000^4^ cells/well and incubated at 37°C with 5% CO_2_. 24 hours later, various concentrations of AvFc, AvFc^lec-^, or CTX were added to target cells along with the Jurkat NFAT-luc FcγRIIIa-expressing cell line (Jur-FcγRIIIa; Promega, Madison, WI) at a ratio of 15:1. FcγRIIIa signaling activates the NFAT transcription factor, inducing the expression of firefly luciferase through an NFAT responsive promoter. After co-culture for 24 hours, firefly luciferase activity was measured using the Britelite Plus Reporter Gene Assay System (Perkin Elmer, Waltham, MA) on a Synergy HT luminometer (BioTeck, Winooski, VT). Jur-FcγRIIIa cells co-cultured with the target cells in the absence of antibody provided no antibody control luciferase production levels, which were subtracted from the actual signals to yield antibody-specific activation, in relative light units (RLUs). Background determined by taking the mean of the target-cell-only wells. Fold induction was calculated using the following equation: Fold induction = (RLU_induced_ -RLU_background_) / (RLU_noAbcontrol_ -RLU_background_).

### ADCC assay with primary human PBMC effector cells

Similar to the reporter assay, plated A549 target cells were pre-incubated with serial dilutions of AvFc or CTX for 30 min at 37°C. PBMCs were added to initiate the ADCC at ratio of 50:1 for AvFc and 25:1 for CTX. After incubation at 37°C incubator for 6 h, cell supernatants were collected and released lactose dehydrogenase was measured and compared to a no-drug control to calculate % target cell lysis. Each sample was tested in triplicate.

### Subcutaneous lung cancer xenograft challenge model

1×10^7^ A549 cells or 5×10^5^ H460 were implanted into the hind-left flanks of 8-week-old female SCID mice. Mice were then randomly organized into three groups and treated with the vehicle (n=10), 25 mg/kg AvFc (n=10), and 25 mg/ kg CTX (n=10). Vehicle treatment consisted of the AvFc formulation buffer (30 mM histidine pH 7.4, 100 mM sucrose, 100 mM NaCl). Treatments were administered i.p. on days 4, 6, 8, 10, 12, and 14 following the formation of palpable lesions. Body weights and tumor volumes were measured every other day after treatment. Animals were euthanized 26 days after A549 challenge and 16 days after H460 challenge.

### A549 lung metastasis model

A549 cells expresing GFP were grown to confluency in growth medium. After harvest, 2×10^6^ cells were injected i.v. into 9-week-old female SCID mice. Mice were then randomly organized into two groups and treated with the vehicle (n=10), and 25 mg/kg AvFc (n=10). Treatments were administered i.p. on days 0, 2, 4, 6, 8, and 10. Following treatment, mice were weighed every other day. Finally, the animals were euthanized on day 28, with the lungs surgically removed and fixed in 10% formalin. To detect GFP signals, each lobe of lungs was separated and GFP signals were detected by Amersham Imager 600 (GE Healthcare Life Sciences, Chicago, IL).

### Immunohistochemistry

Immunohistochemical staining was performed on the cryo-sections of frozen tissues from lung patients undergoing surgery. Staining was performed with the VECTASTAIN® Elite® ABC HRP Kit (Peroxidase, Standard) (Vector Labs, Burlingame, CA). 8 µm tissue sections were placed on positively charged slides (VWR International, Radnor, PA) and air dried. Then, sections were incubated for 10 minutes at room temperature in 3% H_2_O_2_ diluted in methanol and washed with Tris-buffered saline supplemented with 0.01% triton-X100 (TBST). Avidin/biotin blockade was performed using a blocking kit for 15 minutes at room temperature (Abcam, Cambridge, MA), then Fc-receptors were blocked in Fc-blocking solution for 10 minutes at room temperature (BD, San Jose, CA). Sections were further blocked with with 3% goat serum in TBST for 30 minutes at room temperature. To stain the tissue, 0.5 µg/ml of AvFc in TBST supplemented with 1% goat serum was added for 30 minutes at room temperature, followed by a biotinylated anti-human IgG in TBST with 1% goat serum also incubated for 30 minutes at room temperature (Vector Labs, Burlingame, CA). Then, ABC solution was added for 30 minutes at room temperature (Vector Labs, Burlingame, CA) followed by the DAB stain, which was applied following the manufacturer’s protocol (Vector Labs, Burlingame, CA). Sections were washed between each step 3 times with TBST. Counter staining was performed with hematoxylin. Sections were serially dehydrated with 95% ethanol, 100% ethanol, and CitriSolv (Decon lab, King of Prussia, PA). Images were taken using an OLYMPUS CKX41 microscope with UPlanFL 10x/0.30 lens (OLYMPUS, Tokyo, Japan).

### Statistical analyses

Group means and standard errors were derived from the values obtained in three individual replicates, and assays were performed at least twice independently unless otherwise noted. For all data, outliers were determined by statistical analysis using the Grubb’s test (*p* < 0.05) and excluded from further analysis. Statistical significance was analyzed by one-way or two-way analysis of variance (ANOVA) with Bonferroni’s post-hoc test or Wilcoxon matched-pairs signed rank test as indicated in figure legends, using GraphPad Prism 5 (San Diego, CA). Differences were considered statistically significant if *p* < 0.05.

## Data availability

– All data generated during or analyzed during the current study are available from the corresponding author on reasonable request.

## Materials & Correspondence

Correspondence and material requests should be addressed to: Nobuyuki Matoba, University of Louisville School of Medicine, 505 S. Hancock Street, Room 615, Louisville, KY 40202, USA, Tel: (502) 852 8412; Fax: (502) 852 5468; E-Mail

## ACKNOWLEDGEMENTS

We thank J. Calvin Kouokam, Rachel Carry and Sucheta Telang for useful discussion and technical assistance. This work was supported by a NIH grant (R21-CA216447), a U.S. Department of Defense grant (W81XWH-10-2-0082-CLIN2) and a University of Louisville Brown Cancer Center Molecular Target CoBRE grant (as a subproject of NIH NIGMS/ P30-GM106396). M.W.D. was supported by a T32 Environmental Health Sciences Grant (T32-ES11564).

## AUTHOR CONTRIBUTIONS

Y.J.O, M.W.D. and N.M. wrote the manuscript. Y.J.O. performed experiments. M.W.D. and A.R.F. assisted to perform parts of experiments. M.L.M. performed proteomics analysis. Y.J.O., M.W.D., Q.Z., M.L.M, C.B.L. and N.M. analyzed data. N.M. conceived and designed experiments, secured funding, and supervised the study.

