## Supplementary material for "Antitumor activity of a lectibody targeting cancer-associated high-mannose glycans": Figure S1, Figure S2, Table S1

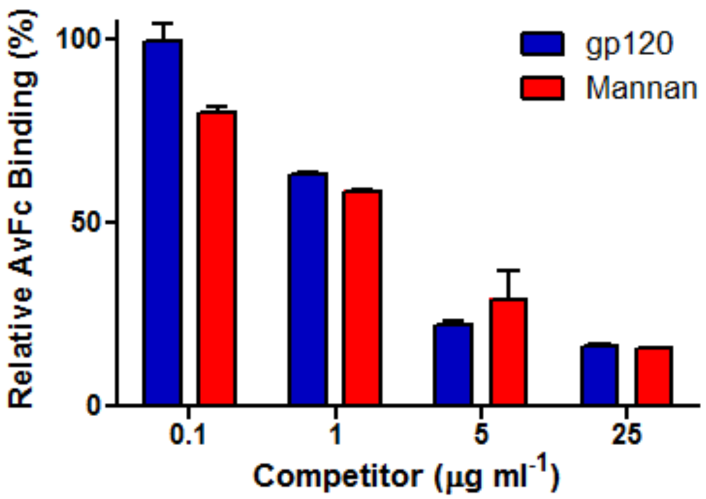


**Figure S1. Inhibition of AvFc binding to cancer cells by HIV-1 gp120 and yeast mannan.** A549 lung cancer cells were incubated with AvFc (1 µg/ml) and various concentrations of HIV-1 envelope glycoprotein gp120 and yeast mannan for 30 minutes at 4ºC. Cells were washed and stained with 10 µg/ml of goat anti-human IgG FITC for 30 minutes at 4ºC. Cells were then washed and analyzed for binding on a FACS Canto II (BD Biosciences) using FACSDiva (BD Biosciences).

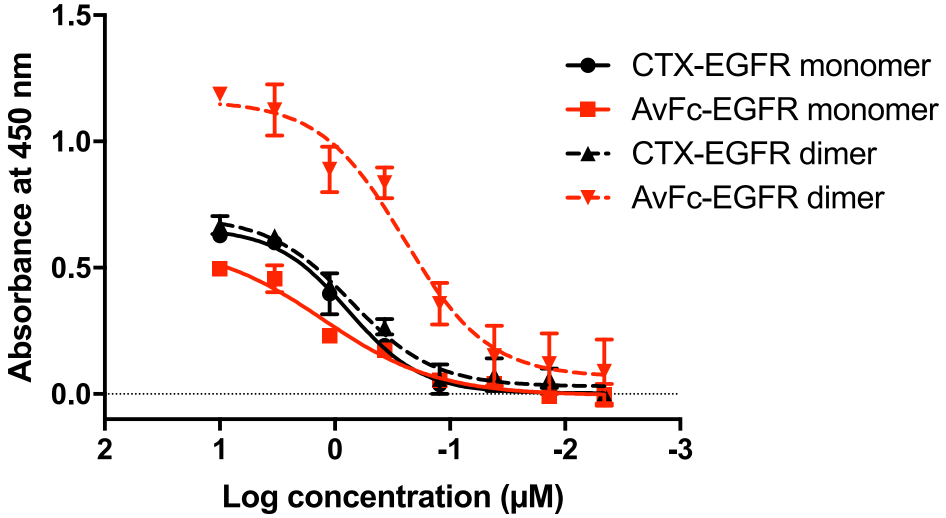


**Figure S2. AvFc and CTX binding to monomeric and dimeric EGFRs.** EGFR monomer and dimer were purified without and with 20 ng/ml of EGF treatments for 10 minutes, respectively, in A549 cells. To maintain a dimer form, EGFR dimer was cross-linked by $BS^3$ linkers (Thermo Fisher Scientific, 21580) and filtered by 200kDa cut-off filter (Advantec, USY-20). Vesicles composed of 1-palmitoyl-2-oleoyl-glycero-3-phosphocholine (POPC) and 1,2-dioleoyl-sn-glycero-3-phospho-L-serine (DOPS) (90:10 in molar ratio). Uniform sized (100 nm) vesicles were prepared by the vesicle extrusion method. EGFR (monomer) or EGFR (dimer) were inserted into vesicles (protein: lipids=2000:1 in molar ratio) in 0.7% of CHAPS added PBS. After dialysis in 2 L of PBS, overnight at 4°C, varying concentrations of EGFR proteo-liposome was applied to an ELISA plate coated with 2 µg/ml of an anti-EGFR antibody (rabbit; Cell Signaling Technology, D38B1). The plate-captured EGFR proteo-liposomes were detected with 13.7 nM of AvFc or Cetuximab (CTX) and an anti-human IgG1-HRP conjugate (SouthernBiotech, 9054-05).

### Table S1. Proteomic analysis of AvFc's binding partners in A549 and H460 cells.

Putative A549 cell-surface binding partners

| Gene Name | Uniprot Accession | Percentage of Total Spectra (%) |
|---|---|---|
| *Laminin subunit beta-1* | sp\|P07942\|LAMB1_HUMAN | 0.4100 |
| *Laminin subunit gamma-1* | sp\|P11047\|LAMC1_HUMAN | 0.3600 |
| *Laminin subunit alpha-5* | sp\|O15230\|LAMA5_HUMAN | 0.3400 |
| *Agrin* | sp\|O00468-3\|AGRIN_HUMAN | 0.2100 |
| *Mucin-5AC* | sp\|P98088\|MUC5A_HUMAN | 0.2100 |
| *Cation-independent mannose-6-phosphate receptor* | sp\|P11717\|MPRI_HUMAN | 0.2000 |
| *Epidermal growth factor receptor* | sp\|P00533\|EGFR_HUMAN | 0.1900 |
| *Isoform 6 of Myoferlin* | sp\|Q9NZM1-6\|MYOF_HUMAN | 0.1700 |
| *Polycystin-2* | sp\|Q13563-3\|PKD2_HUMAN | 0.1700 |
| *Transferrin receptor protein 1* | sp\|P02786\|TFR1_HUMAN | 0.1600 |
| *Integrin alpha-V* | sp\|P06756-2\|ITAV_HUMAN | 0.1500 |
| *Erlin-2* | sp\|O94905\|ERLN2_HUMAN | 0.1500 |
| *Integrin alpha-3* | sp\|P26006\|ITA3_HUMAN | 0.1500 |
| *Mucin-5B* | sp\|Q9HC84\|MUC5B_HUMAN | 0.1400 |
| *Tetratricopeptide repeat protein 13* | sp\|Q8NBP0\|TTC13_HUMAN | 0.1300 |
| *CD109 antigen* | sp\|Q6YHK3\|CD109_HUMAN | 0.1300 |
| *Tetratricopeptide repeat protein 17* | sp\|Q96AE7-2\|TTC17_HUMAN | 0.1300 |
| *Plexin-B2* | sp\|O15031\|PLXB2_HUMAN | 0.1200 |
| *Protocadherin Fat 1* | sp\|Q14517\|FAT1_HUMAN | 0.1200 |
| *Sortilin-related receptor* | sp\|Q92673\|SORL_HUMAN | 0.1200 |
| *Transmembrane protein 2* | sp\|Q9UHN6-2\|TMEM2_HUMAN | 0.1100 |
| *Integrin beta-5* | sp\|P18084\|ITB5_HUMAN | 0.1100 |
| *Spectrin beta chain, non-erythrocytic 1* | sp\|Q01082\|SPTB2_HUMAN | 0.1100 |
| *Erlin-1* | sp\|O75477\|ERLN1_HUMAN | 0.1100 |
| *Integrin alpha-2* | sp\|P17301\|ITA2_HUMAN | 0.1100 |
| *Transmembrane and TPR repeat-containing protein 3* | sp\|Q6ZXV5-2\|TMTC3_HUMAN | 0.1100 |
| *Contactin-associated protein-like 3* | sp\|Q9BZ76\|CNTP3_HUMAN | 0.1100 |
| *Disintegrin and metalloproteinase domain-containing protein 9* | sp\|Q13443\|ADAM9_HUMAN | 0.1100 |
| *Transmembrane protein 106B* | sp\|Q9NUM4\|T106B_HUMAN | 0.0980 |
| *Insulin-like growth factor 1 receptor* | sp\|P08069\|IGF1R_HUMAN | 0.0910 |
| *Metal transporter CNNM4* | sp\|Q6P4Q7\|CNNM4_HUMAN | 0.0910 |
| *Adhesion G protein-coupled receptor L2* | sp\|O95490-2\|AGRL2_HUMAN | 0.0910 |
| *T-complex protein 1 subunit theta* | sp\|P50990\|TCPQ_HUMAN | 0.0910 |
| *Attractin* | sp\|O75882\|ATRN_HUMAN | 0.0840 |
| *Tropomyosin alpha-3 chain* | sp\|P06753-2\|TPM3_HUMAN | 0.0700 |
| *Multidrug resistance-associated protein 1* | sp\|P33527-3\|MRP1_HUMAN | 0.0700 |
| *Galectin-3-binding protein* | sp\|Q08380\|LG3BP_HUMAN | 0.0700 |
| *Serpin H1* | sp\|P50454\|SERPH_HUMAN | 0.0630 |
| *Prolow-density lipoprotein receptor-related protein 1* | sp\|Q07954\|LRP1_HUMAN | 0.0560 |
| *Exportin-1* | sp\|O14980\|XPO1_HUMAN | 0.0560 |
| *Disintegrin and metalloproteinase domain-containing protein 17* | sp\|P78536\|ADA17_HUMAN | 0.0560 |
| *Importin subunit beta-1* | sp\|Q14974\|IMB1_HUMAN | 0.0560 |
| *Transmembrane protein 131* | sp\|Q92545\|TM131_HUMAN | 0.0560 |
| *Laminin subunit beta-2* | sp\|P55268\|LAMB2_HUMAN | 0.0560 |
| *Scavenger receptor class B member 1* | sp\|Q8WTV0-2\|SCRB1_HUMAN | 0.0560 |
| *Sodium/potassium-transporting ATPase subunit beta-1* | sp\|P05026-2\|AT1B1_HUMAN | 0.0560 |
| *Integrin alpha-5* | sp\|P08648\|ITA5_HUMAN | 0.0350 |
| *Cation-dependent mannose-6-phosphate receptor* | sp\|P20645\|MPRD_HUMAN | 0.0280 |
| *Solute carrier family 12 member 7* | sp\|Q9Y666\|S12A7_HUMAN | 0.0210 |
| *Integrin alpha-1* | sp\|P56199\|ITA1_HUMAN | 0.0210 |
| *Cell cycle control protein 50A* | sp\|Q9NV96\|CC50A_HUMAN | 0.0210 |
| *Magnesium transporter protein 1* | sp\|Q9H0U3\|MAGT1_HUMAN | 0.0210 |
| *Neutral amino acid transporter B (0)* | sp\|Q15758\|AAAT_HUMAN | 0.0140 |
| *Endothelial protein C receptor* | sp\|Q9UNN8\|EPCR_HUMAN | 0.0140 |
| *Solute carrier family 2, facilitated glucose transporter member 1* | sp\|P11166\|GTR1_HUMAN | 0.0070 |

Putative H460 cell-surface binding partners

| | | |
|---|---|---|
| *Laminin subunit gamma-1* | sp\|P11047\|LAMC1_HUMAN | 0.8300 |
| *Cation-independent mannose-6-phosphate receptor* | sp\|P11717\|MPRI_HUMAN | 0.4400 |
| *Laminin subunit alpha-1* | sp\|P25391\|LAMA1_HUMAN | 0.3500 |

| | | |
|---|---|---|
| *Laminin subunit beta-1* | sp\|P07942\|LAMB1_HUMAN | 0.3100 |
| *Laminin subunit alpha-5* | sp\|O15230\|LAMA5_HUMAN | 0.2300 |
| *ATP-binding cassette sub-family A member 2* | sp\|Q9BZC7-3\|ABCA2_HUMAN | 0.2100 |
| *Fibronectin* | sp\|P02751-17\|FINC_HUMAN | 0.2100 |
| *Transferrin receptor protein 1* | sp\|P02786\|TFR1_HUMAN | 0.2100 |
| *Galectin-3-binding protein* | sp\|Q08380\|LG3BP_HUMAN | 0.1700 |
| *Laminin subunit beta-2* | sp\|P55268\|LAMB2_HUMAN | 0.1600 |
| *Scavenger receptor class B member 1* | sp\|Q8WTV0-2\|SCRB1_HUMAN | 0.1500 |
| *Agrin* | sp\|O00468-3\|AGRIN_HUMAN | 0.1500 |
| *Plexin-B2* | sp\|O15031\|PLXB2_HUMAN | 0.1300 |
| *Contactin-associated protein 1* | sp\|P78357\|CNTP1_HUMAN | 0.1300 |
| *Integrin alpha-1* | sp\|P56199\|ITA1_HUMAN | 0.1300 |
| *CD109 antigen* | sp\|Q6YHK3-4\|CD109_HUMAN | 0.1300 |
| *Filamin-A* | sp\|P21333-2\|FLNA_HUMAN | 0.1300 |
| *Integrin alpha-V* | sp\|P06756-2\|ITAV_HUMAN | 0.1300 |
| *Integrin beta-5* | sp\|P18084\|ITB5_HUMAN | 0.1200 |
| *Myoferlin* | sp\|Q9NZM1-2\|MYOF_HUMAN | 0.1100 |
| *Tetratricopeptide repeat protein 13* | sp\|Q8NBP0-2\|TTC13_HUMAN | 0.1100 |
| *Transmembrane protein 106B* | sp\|Q9NUM4\|T106B_HUMAN | 0.1100 |
| *Neural cell adhesion molecule L1* | sp\|P32004-2\|L1CAM_HUMAN | 0.0990 |
| *Scavenger receptor class A member 5* | sp\|Q6ZMJ2\|SCAR5_HUMAN | 0.0990 |
| *Disintegrin and metalloproteinase domain-containing protein 9* | sp\|Q13443\|ADAM9_HUMAN | 0.0910 |
| *4F2 cell-surface antigen heavy chain* | sp\|P08195-2\|4F2_HUMAN | 0.0910 |
| *Metal transporter CNNM4* | sp\|Q6P4Q7\|CNNM4_HUMAN | 0.0830 |
| *Secretory carrier-associated membrane protein 3* | sp\|O14828\|SCAM3_HUMAN | 0.0830 |
| *Renin receptor* | sp\|O75787-2\|RENR_HUMAN | 0.0660 |
| *Niemann-Pick C1 protein* | sp\|O15118\|NPC1_HUMAN | 0.0660 |
| *Synaptonemal complex protein SC65* | sp\|Q92791\|SC65_HUMAN | 0.0660 |
| *Nidogen-2* | sp\|Q14112-2\|NID2_HUMAN | 0.0660 |
| *Urokinase plasminogen activator surface receptor* | sp\|Q03405-2\|UPAR_HUMAN | 0.0660 |
| *Epidermal growth factor receptor* | sp\|P00533\|EGFR_HUMAN | 0.0580 |
| *Sortilin-related receptor* | sp\|Q92673\|SORL_HUMAN | 0.0580 |
| *Serpin H1* | sp\|P50454\|SERPH_HUMAN | 0.0580 |
| *Attractin* | sp\|O75882-2\|ATRN_HUMAN | 0.0580 |
| *Protocadherin Fat 1* | sp\|Q14517\|FAT1_HUMAN | 0.0580 |
| *Neutral amino acid transporter B (0)* | sp\|Q15758\|AAAT_HUMAN | 0.0500 |
| *Integrin alpha-5* | sp\|P08648\|ITA5_HUMAN | 0.0410 |
| *Cell cycle control protein 50A* | sp\|Q9NV96\|CC50A_HUMAN | 0.0330 |
| *Solute carrier family 2, facilitated glucose transporter member 1* | sp\|P11166\|GTR1_HUMAN | 0.0330 |
| *Endothelial protein C receptor* | sp\|Q9UNN8\|EPCR_HUMAN | 0.0330 |
| *Insulin-like growth factor 1 receptor* | sp\|P08069\|IGF1R_HUMAN | 0.0330 |
| *Integrin alpha-3* | sp\|P26006\|ITA3_HUMAN | 0.0250 |
| *Cation-dependent mannose-6-phosphate receptor* | sp\|P20645\|MPRD_HUMAN | 0.0250 |
| *Prolow-density lipoprotein receptor-related protein 1* | sp\|Q07954\|LRP1_HUMAN | 0.0170 |
| *Magnesium transporter protein 1* | sp\|Q9H0U3\|MAGT1_HUMAN | 0.0170 |
| *Solute carrier family 12 member 7* | sp\|Q9Y666\|S12A7_HUMAN | 0.0083 |
| *Integrin alpha-2* | sp\|P17301\|ITA2_HUMAN | 0.0083 |

Co-immunoprecipitation with AvFc covalently bound to agarose beads was used to capture potential binding partners in whole-cell lysates. These proteins were then identified using mass spectrometry. The top 100 hits were then narrowed down using Gene Ontology terms as well as literature searches so as to include only transmembrane receptors, transporters, and/or adhesion molecules which may be present on the cell surface and thus which may have increased incidence of high-mannose glycans. Putative binding partners are ranked by their relative abundance within each individual analysis.
